# Defining the developmental program leading to meiosis in maize

**DOI:** 10.1101/434993

**Authors:** Brad Nelms, Virginia Walbot

## Abstract

In multicellular organisms, the entry into meiosis is a complex process characterized by increasing levels of meiotic specialization. We used single-cell RNA-sequencing to reconstruct the developmental program into meiosis in maize. We observed a smooth continuum of expression stages leading up to meiosis, followed by a sharp reorganization of the transcriptome in early meiotic prophase. This latter transcriptional shift was dramatic, with 26.7% of expressed genes changing by 2 fold or more, and occurred just prior to a proposed cell cycle checkpoint. Changes in cell physiology accompanied the nuclear events of meiosis, including a decrease in protein translation capacity and increase in membrane-bound organelles. We further identified differences in gene expression between the mitotic and meiotic cell cycles. Our results uncover a multi-step pathway into meiosis and highlight the power of single cell RNA-seq to define developmental transitions.

---

Meiosis is a pivotal event in the life cycle of sexually reproducing organisms, reducing the chromosome number in half and creating new allele combinations through recombination. The mechanisms that regulate meiotic entry in plants are not well understood^1,2^, but are of great importance to crop breeding and agricultural yield^3^.

In maize anthers, archesporial (AR) cells are the first cell type in the lineage dedicated to a meiotic fate, acting as the developmental switch between somatic and germinal growth^4^. After a ~3 day period of transit amplifying mitotic divisions, AR cells become pollen mother cells (PMCs) and then enter meiotic prophase. There are substantial changes in cell morphology^4^ and gene expression^5–8^ during pre-meiotic and early meiotic development, but the cellular intermediates that arise during this process are not well-defined.

Here, we applied single-cell RNA-sequencing (scRNA-seq) to characterize the developmental program leading up to meiosis in maize with high resolution. We introduce a quantitative framework, ‘pseudotime velocity’, to infer developmental transitions based on periods of relatively rapid gene expression change, and apply this framework to identify intermediates from our data. These results provide a road-map for reconstructing plant developmental pathways with scRNA-seq.

## Single-cell RNA-seq of pre-meiotic and early meiotic cells

Maize male germinal cells form within immature anthers, centrally located in each of four anther lobes (Fig. 1A). Anthers expand in size predictably during early development; consequently, anther length can be used as a reliable, continuous staging system that is correlated with both developmental events and organ age^4^. We established methods to isolate single pre-meiotic and meiotic cells from maize anthers for scRNA-seq (Fig. 1B). Cells in the germinal lineage are 2-4 times larger in diameter than all other anther cells, permitting their identification after tissue dissociation (Fig. 1B, right). To minimize expression changes during sample handling, we optimized protoplasting conditions and established a rapid manual cell isolation protocol (see Methods). In the final procedure, isolated cells were obtained and frozen within 1-2 hours of plant harvest. After cell lysis, all single-cell samples were divided into two tubes and processed independently as split-cell technical replicates.

**Fig. 1.**
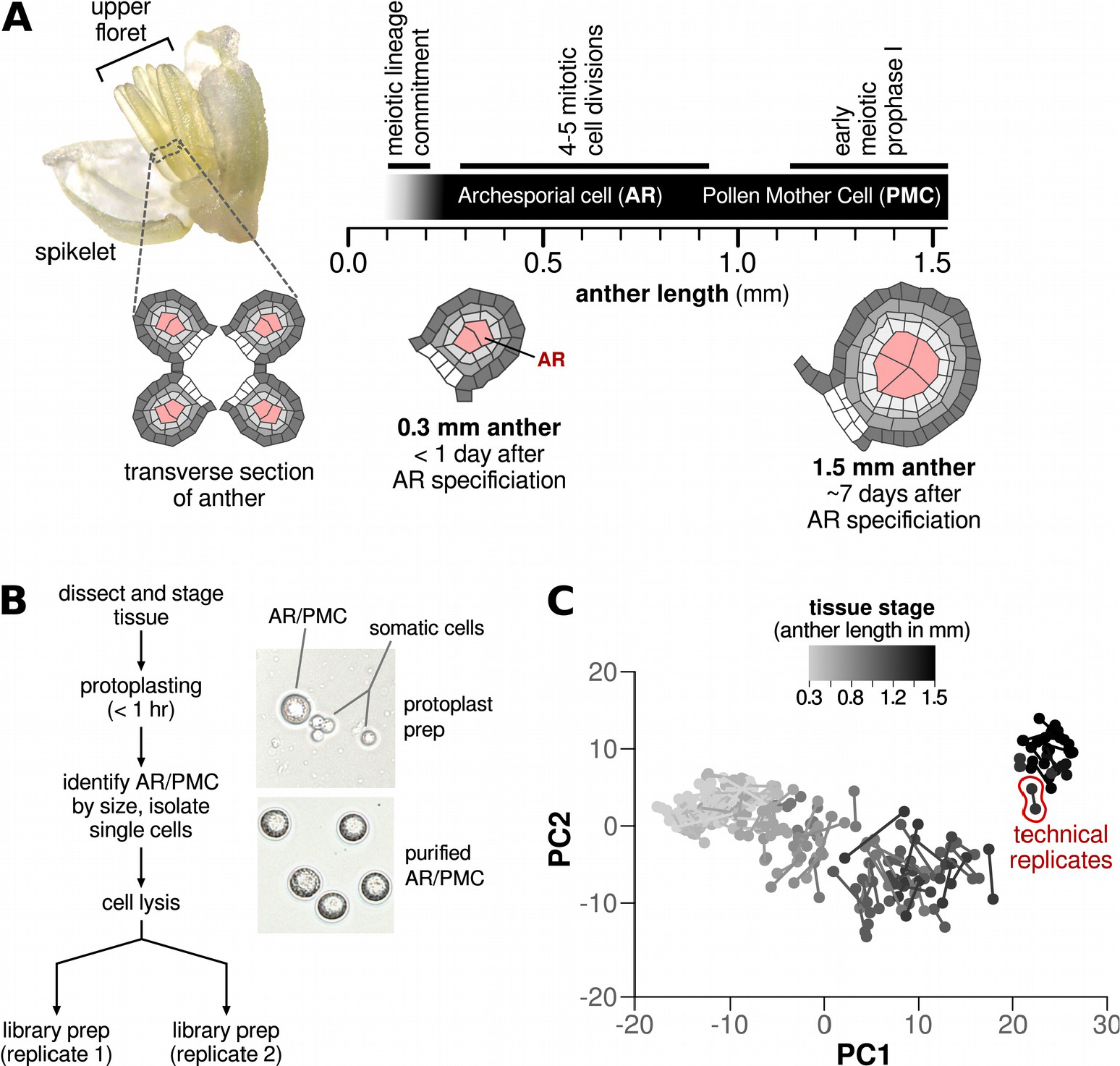
Single-cell RNA-seq of pre-meiotic and early meiotic cells in maize. (**A**) Schematic of early anther development. Single cells were isolated from anthers between 0.3 and 1.5 mm in length; anther length is a reliable proxy for tissue stage during this period of development. (**B**) Flow chart of cell isolation and single-cell library preparation. AR/PMCs were identified based on their large size; technical replicates were prepared by splitting single-cell contents immediately after cell lysis. (**C**) Principal component analysis (PCA) shows that major axes of variation in the dataset closely reproduce tissue stage. Replicate samples were considered independently then connected with a line for visualization.

Using this protocol, we isolated pre-meiotic and early meiotic cells from 24 plants, covering a week of development from the day after AR cell specification to the early zygotene stage of meiotic prophase I. Starting material was staged by anther length, and anther stages were densely sampled from throughout this period (Fig. S1 and Table S1). High quality reads were obtained from 144 cells (Fig. S2), with a mean of 101,245 distinct transcripts per cell. The number of transcripts detected is substantially higher than typically observed for single-cell data in animals, likely reflecting the relatively large amount of RNA present in cells from the maize germinal lineage (~150 pg of total RNA per cell compared to 5-20 pg for most animal cell types). Technical replicates provided reproducible transcript counts (Fig. 1C and S3; median *r*^*2*^ = 0.92), demonstrating the quality of this dataset. Furthermore, the major axes of variation across the data were closely correlated with tissue stage (Fig. 1C). A set of 375 genes (Table S2; 2.1% of the 12,902 ‘expressed’ genes) formed clusters indicative of stage in the cell cycle (Fig. S4). To focus on developmental events, these cell cycle-regulated genes were excluded from all analyses unless specified otherwise.

## Dynamics of early germinal cell differentiation

To identify intermediate cellular stages from our data, we initially applied cell clustering to group cells into distinct bins. However, clustering removed valuable information about developmental dynamics: some clusters had much sharper boundaries than others, and many had internal structure not well-captured by discrete cell groupings. As a result, we moved to a continuous framework to assess cell-to-cell variation and developed a statistic, ‘pseudotime velocity’, to quantify variation in the rate of gene expression change over time.

First, we applied dimensionality reduction to our data to determine pseudotime^9,10^, a single variable that captured much of the relative gene expression difference between cells as well as their inferred developmental order (Fig. 2A). Pseudotime estimates were reproducible between split-cell technical replicates (*r*^*2*^ = 0.97; Fig. 2A, inset). We then calculated pseudotime velocity as the local linear slope in pseudotime between cells (Fig. 2B); this metric was inherently noisy, but repeated bootstrap sampling dampened this noise and allowed for the quantification of uncertainty in the velocity estimates (Fig. S5).

**Fig. 2.**
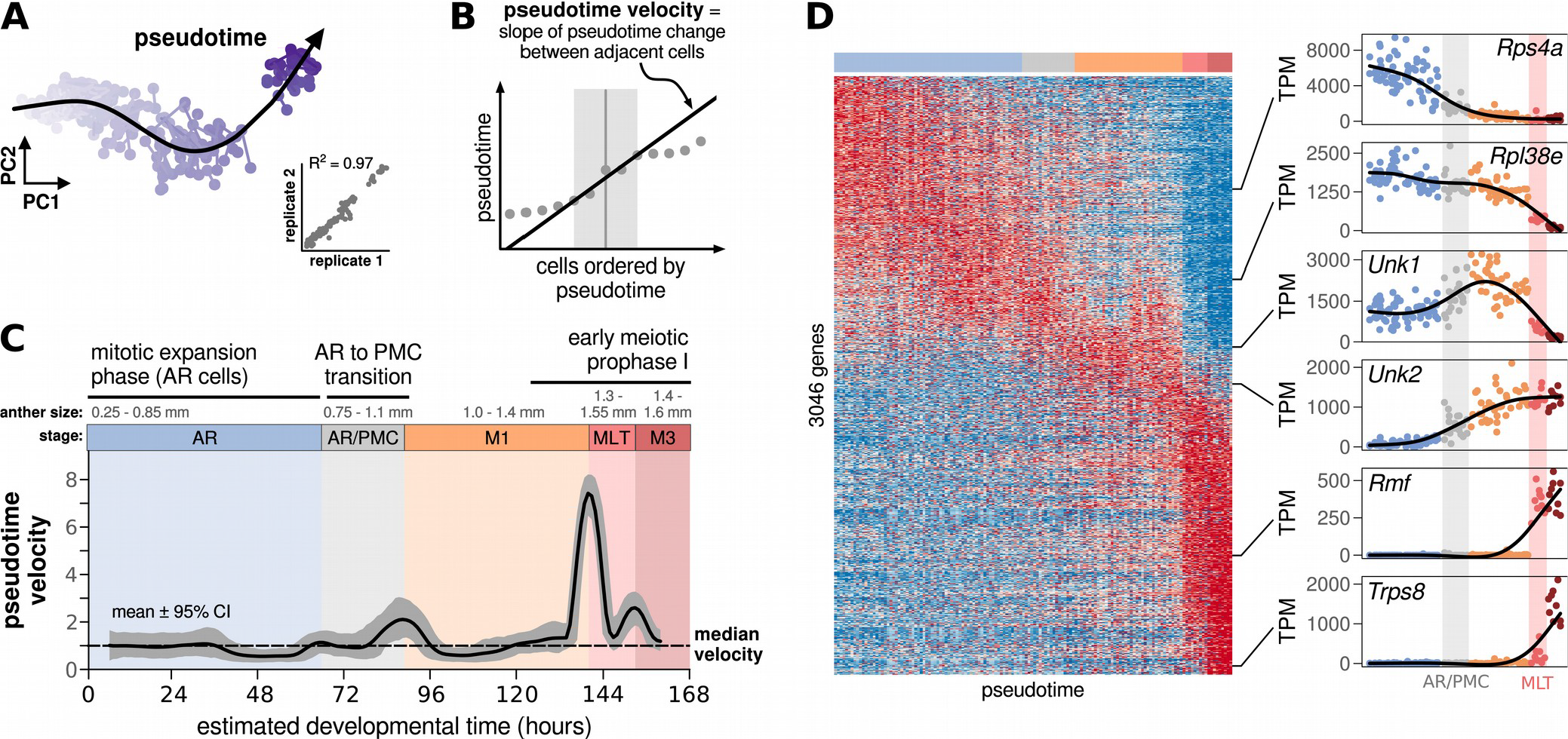
Germinal differentiation is characterized by both gradual and discrete cell state transitions. (**A**) Pseudotime was calculated by fitting a principal curve to the first ten principal components. Inset, pseudotime estimates were reproducible between technical replicates. (**B**) Pseudotime velocity was calculated as the local slope of pseudotime with a 10 sample window; results were consistent with a range of window sizes from 2-15 samples. (**C**) Pseudotime velocity as a function of estimated developmental time. Units are relative to median velocity and the gray outline denotes the 95% confidence interval. Four peaks with relatively rapid pseudotime change were selected as stage boundaries, highlighted by colored boxes. AR, archesporial cell; PMC, Pollen Mother Cell; M1, meiotic stage 1; MLT, mid-leptotene transition (see below); M3, meiotic stage 3. (**D**) Heatmap of gene expression for all differentially regulated genes. Color ranges from blue (minimum TPM) to red (maximum TPM) for each gene. There are several waves of gene expression during early germinal development, then two sharp shifts during early meiotic prophase I. TPM, transcripts per million; *Unk1*, unknown gene 1 (Zm00001d027037); *Unk2*, unknown gene 2 (Zm00001d013377).

Pseudotime velocity varied substantially during the time-course, with several noticeable peaks in the velocity curve (Fig. 2C). These peaks agreed with cluster borders defined by the consensus of 8 single-cell clustering methods (Fig. S6A), and, remarkably, outperformed all clustering methods at accurately grouping split-cell technical replicates (Fig. S6B). We defined the four most prominent peaks as stage boundaries (colored boxes in Fig. 2C), providing a refined staging system for early germinal differentiation in maize. The stages were associated with specific anther lengths, making it possible to connect each to the established developmental progression; there were stages consistent with AR (transit-amplifying) cells, the AR-to-PMC transition, early meiosis (M1), and two stages during early meiotic prophase I (MLT and M3, see below). Thus, pseudotime velocity captured biologically meaningful stages separated by short periods of relatively rapid gene expression change.

Pseudotime velocity also preserved many details obscured by clustering. For example, there were two periods with a statistically significant *decrease* in velocity compared to the median. One of these occurred in the middle of the AR stage (0.5 – 0.8 mm anthers), and may represent a pause in differentiation while the AR cells expanded mitotically prior to meiosis. Second, pseudotime velocity provided a means to quantify the relative strength of each stage boundary. The first velocity peak in early meiotic prophase was substantially larger than all others, reaching a 7.4-fold higher velocity than the median (95% confidence interval [CI] = 6.5, 8.2). In contrast, the velocity remained under 2-times the median throughout the entire AR period (*p* < 0.05, permutation test), despite the fact that one third of all pseudotime change occurred during this period. To investigate the underlying features leading to the observed differences in velocity, we visualized the expression of all genes that were significantly correlated with pseudotime (3046 genes; Table S3) in a heatmap (Fig. 2D). The early stages showed smooth, continuous changes with successive waves of gene expression, while the final two stages were characterized by step-like discrete shifts in expression. Thus we find evidence for varying periods of continuous and discrete modes of differentiation in this single lineage.

## A major shift in gene expression during mid-leptotene

The final two stages we identified (MLT and M3 in Fig. 2C) occurred in 1.3 to 1.5 mm anthers, when the PMCs were in early prophase I of meiosis. Maize meiocytes progress through a series of classic cytological stages during this time period^11^ (Fig. 3A, top), but these stages cannot be unambiguously identified by anther length. To relate the gene expression shifts we observed to the established cytological progression, we directly compared gene expression to chromosome cytology in 1.1 to 1.6 mm anthers isolated from 47 individual florets. One of three synchronously developing anthers from each floret was fixed for cytology, while meiotic cells were harvested from the other two anthers for quantitative PCR (qPCR) of five stage markers selected from the scRNA-seq data (Fig. S7).

**Fig. 3.**
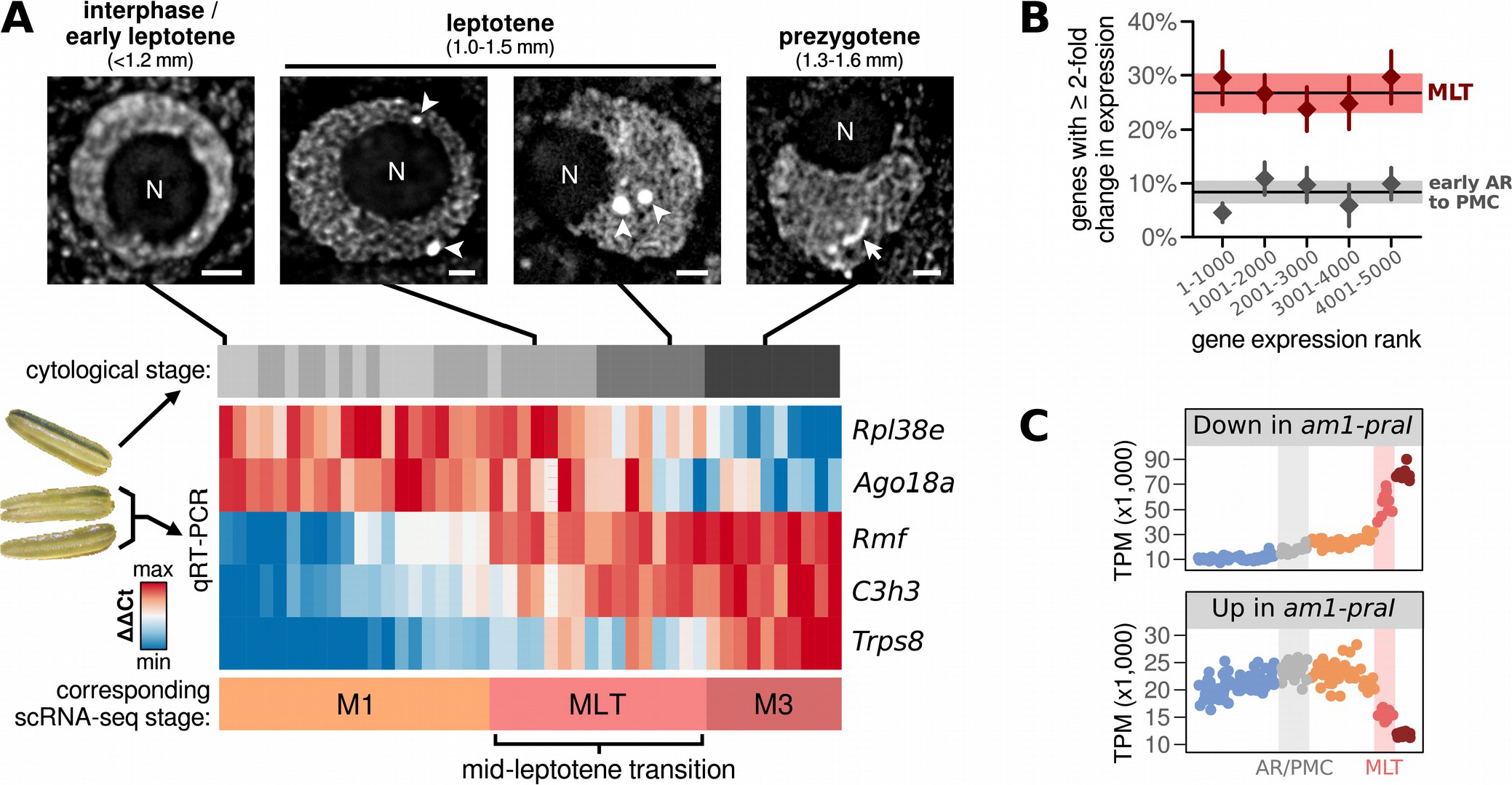
Reorganization of the transcriptome in mid-leptotene. (**A**) Comparison of gene expression stages with cytology in 1.1 to 1.6 mm anthers. (top) Hoescht staining of meiotic nuclei from anthers at each cytological stage. Anthers were divided into four cytological stages based on the presence or absence of condensed chromosome throughout the nuclear volume, the location of the nucleolus at the center or periphery of the nucleus, and the spherical or elongated morphology of knobs. N, nucleolus; arrowhead, spherical knobs; arrow, elongated knob. Scale bar, 2 μm. (bottom) Quantitative RT-PCR on isolated meiocytes for 5 marker genes selected from the single-cell data. (**B**) Estimated percentage of genes with a ≥2 fold change in expression during the mid-leptotene transition (MLT) and from early AR (pseudotime − 10) to the M1 stage. The black horizontal lines and shaded area show the mean +/− 95% CI for the estimated percentage of genes that change ≥2 fold when calculated with the top 5000 expressed genes; triangles show the estimates for subsets of these genes grouped by overall expression level. (**C**) Cumulative expression of transcripts established^14^ to be up- or down- regulated in *am1-praI* mutant anthers.

We found that expression by qPCR reproduced the expression stages observed by single-cell RNA-seq (Fig. 3A, heatmap with corresponding scRNA-seq stages labeled): there was an early stage characterized by high expression of *Rpl38e* and *Ago18a* and low expression of *Rmf*, *C3h3*, and *Trps8* (comparable to stage M1 in Fig. 2C), an intermediate stage with increased expression of the markers *Rmf* and *C3h3* (MLT in Fig. 2C), and a final stage characterized by decreased expression of *Rpl38e* and *Ago18a* and increased expression of *Trps8* (M3 in Fig. 2C). By comparing these expression results with cytological staging, we found that the middle stage begins in mid-leptotene (Fig. 3A), after chromatin condensation into thin fibers throughout the nuclear volume, and lasts until the prezygotene stage^11^, when condensed heterochromatic structures called knobs elongate. We defined this stage as the ‘mid-leptotene transition’ (MLT) to denote that this expression change occurs in the middle of leptotene. The MLT cannot be uniquely identified by chromosome cytology or anther length, but the expression markers defined here could be used to identify this stage in future work. We estimated that the MLT lasts approximately 24 hours, equivalent to half the duration of leptotene^12^.

The extent of expression change during the MLT was enormous, with a 2-fold or greater change in transcript abundance for 26.7% of expressed genes (95% confidence interval [CI] = 23.1%, 30.3%; Fig. 3B). By comparison, we observed a 2-fold change in only 8.4% of transcripts between the earliest AR cells and the start of meiotic prophase I (CI = 6.2%, 10.5%) – a 5-day development period that included the mitosis-to-meiosis transition. These findings were consistent for genes at different thresholds of overall transcript abundance (Fig. 3B) and were not sensitive to normalization (Fig. S8). The shift in gene expression during the MLT cannot be explained solely by the downregulation of pre-existing transcripts, as there was a large expression increase for a substantial number of genes (including 33 that were induced by 10 fold or more; Table S2). In addition, there is no indication of a change in overall mRNA content based on the number of transcripts detected per cell or the proportion of transcripts relative to spike-in controls (Fig. S9).

It is intriguing that there was a major shift in gene expression leading up to the prezygotene stage. Prezygotene has been proposed to be a cell cycle checkpoint in plants because several mutations result in a phenotype where meiocytes are stalled at this stage^13^. To determine whether mutants that stall in prezygotene were defective in expressing genes characteristic of the MLT, we compared the expression changes during the MLT to the expression changes observed in *am1-praI* mutant plants^14^, which are uniformly stalled in prezygotene^13^. We found that genes downregulated in *am1-praI* mutant anthers increased in expression during the MLT (Fig. 3C), increasing ~2 fold at the start of meiosis and then another 5 fold during the MLT. Conversely, genes that were upregulated in *am1-praI* anthers showed a decrease in expression during the MLT (Fig. 3C, bottom). Therefore, failure to progress through the MLT explains much of the expression changes observed in the *am1-praI* mutant. Further work will be required to distinguish whether the *am1-praI* mutant is simply delayed in meiotic progression or stalled specifically before or during the MLT.

## Coordination between nuclear and cytosolic events

To determine which genes and pathways were regulated during early germinal differentiation, we grouped genes with similar expression patterns by unsupervised clustering and looked for enriched pathways in each cluster using AgriGO^15^ (Fig. 4A and Table S4). This analysis highlighted broad changes in both metabolism and cell biology during early germinal differentiation. There was a significant shift in the proportion of transcripts annotated to specific cellular organelles (Fig 4A and Fig. S10), including an upregulation of genes linked to the membrane-bound organelles mitochondria, ER, and Golgi during the MLT (clusters 5 + 6). These changes suggest that complex cellular remodeling is coordinated with the nuclear changes that occur during early meiosis, consistent with results from baker’s yeast^16^.

**Fig. 4.**
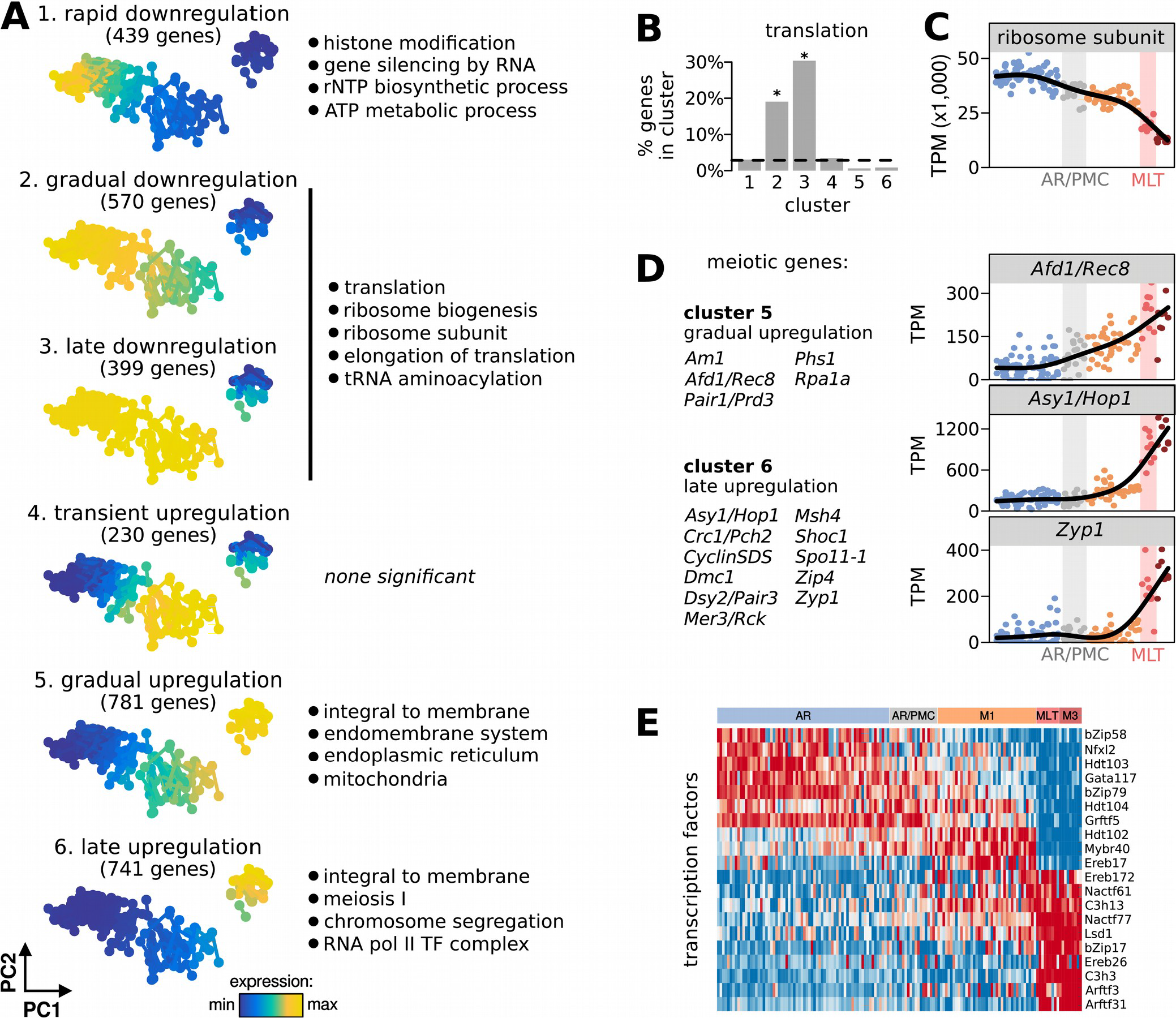
A global view of transcriptional regulation during germinal differentiation. (**A**) PCA plots showing the mean expression of genes in six co-regulated clusters. Select enriched Gene Ontology (GO) terms are listed to the right. (**B**) Percentage of genes in each cluster annotated with the GO term ‘translation’. (**C**) Cumulative expression of transcripts that encode protein subunits of the cytolosic ribosome. (**D**) Expression of genes with established roles in meiotic prophase I. (**E**) Heatmap of the transcript levels for the 20 most differentially expressed TFs.

One striking finding was a dramatic reduction in the protein translation machinery during germinal differentiation. Genes associated with the GO term ‘translation’ composed 20% and 30% of genes in the two major downregulated gene clusters (clusters 2 and 3, respectively; Fig. 4B). These downregulated genes were involved in nearly every aspect of protein translation, including ribosome biogenesis, translation elongation, tRNA metabolism, and structural subunits of the cytosolic ribosome. The extent of this downregulation was large; for instance, the total expression of all ribosomal subunits decreased by 3.4-fold over the time-course (Fig. 4C), with the sharpest decline (2.4-fold) occurring during the MLT. These findings substantiate classic electron microscopy studies^17^, which showed that the number of cytosolic ribosomes decreased by 10 fold or more during early meiotic prophase I in a variety of plant species. Our data show that this ribosome loss is accompanied by a broad shift in overall gene expression. Unexpectedly, the decrease in translation was not accompanied by a repression of transcription, as many transcripts were sharply upregulated during the MLT (Fig. 2D).

Transcripts from genes required for core meiotic functions, such as synapsis and recombination, were abundant in both upregulated gene clusters (clusters 5 and 6; Fig. 4C). There was a loose correlation between when each meiotic gene was upregulated and the established timing of the encoded protein function. For example, the meiotic cohesin subunit *Afd1/Rec8* was upregulated before the axial element protein *Asy1/Hop1*, followed by the transverse filament protein *Zyp1* (Fig. 4C). Because there was tight temporal regulation of transcription for many recombination and synapsis genes, this dataset provides a valuable source of information for the discovery of additional meiotic genes. To establish a revised set of meiotic candidate genes, we took the overlap between genes up-regulated during the MLT (clusters 5 and 6) with a previous list of genes differentially expressed in meiotic cells relative to the surrounding soma^5^. This combination of spatial and temporal information highlighted 374 high priority meiotic candidate genes (Table S3) that are 16-fold enriched for genes known to act in meiotic recombination and synapsis^2^ (*p* = 2.9 × 10^−14^; Fisher’s exact test).

We also observed dynamic expression patterns for many genes families that play key roles in regulating gene expression and cell fate decisions. Sixty five transcription factors (TFs) were differentially regulated during the time-course (Table S3), with expression peaks in each of the major transcriptional stages (Fig. 4E). These include homologs of genes with male-sterility phenotypes in other plant species. For instance, homologs of *C3h3* and *C3h13* are both required for male fertility in *Arabidopsis*, and have been proposed to induce cell wall remodeling during early meiosis^18^; both were upregulated during the MLT. In addition, we observed differential expression for 8 of 18 Argonaute family transcripts, a substantial enrichment from the expected genomic background of 2 (*p* < 0.001; Fisher’s exact test). Argonautes bind diverse classes of small RNA and mediate gene silencing. The dynamic regulation of many Argonaute family members during meiotic differentiation may facilitate the specialized functions of copiously abundant and diverse small RNAs unique to flowers, such as the phasiRNAs^19^.

## Cell cycle regulation in mitosis and meiosis

The asynchronous mitotic divisions during the AR stage complicate efforts to understand gene regulation during this period using bulk genomic approaches. Single-cell RNA-seq provides an opportunity to deconvolve this heterogeneity. We identified several gene clusters within the dataset that appeared to be associated with specific phases of the cell cycle (Fig. 5A, Fig. S4, and Table S2), and inferred the cell cycle phase of each cell based on the expression of these genes (Fig. 5B). The proportion of cells predicted to be in S-phase at a given anther length (Fig. 5C) were in agreement with prior estimates^4^, but the single-cell RNA-seq data also provide a more complete picture because many cell cycle stages are identifiable simultaneously. At the peak cycling stage (0.5 mm anthers), nearly 100% of cells were at some phase of the cell cycle other than G1. This does not support a model where a subset of cells might serve as a stem-cell population that divides asymmetrically to populate the anther lobe with pre-meiotic cells, but rather suggests coordinated tissue-level control of cell cycle potential with all AR cells contributing to the mitotic divisions.

**Fig. 5.**
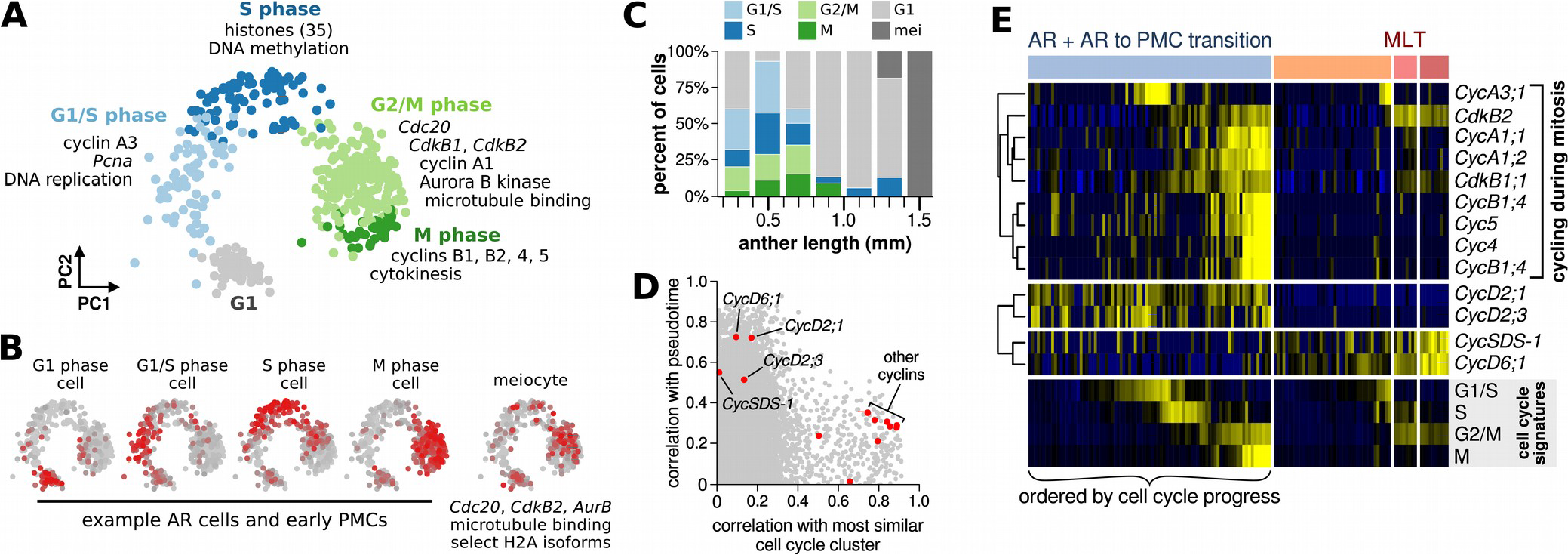
Cell cycle-regulated gene expression in mitosis and meiosis. (**A**) PCA plot of cell cycle-regulated genes. Each point represents a gene, colored based on its assigned cell cycle cluster. (**B**) Expression of cell cycle-regulated genes in representative cells at difference phases in the cell cycle. (**C**) Estimated proportion of cells assigned to each cell cycle stage. (**D**) Correlation of all genes with the most similar mitotic cell cycle cluster and with pseudotime. Cyclins are highlighted in red. (**E**) Heatmap of gene expression for cyclins and cyclin-dependent kinases, grouped by whether they are expressed (top) at specific phases of the cell cycle, (middle) constitutively during the mitotic expansion phase, and (bottom) selectively during meiotic prophase I.

We next compared the expression of the ‘cell cycle-regulated’ genes in mitosis and meiosis. For AR cells and PMCs that had not yet entered meiotic prophase I, these genes were expressed in a cyclical pattern with expression of genes associated with specific cell cycle phases (Fig. 5B). Meiotic cells, in contrast, co-expressed genes associated with multiple mitotic cell cycle phases (Fig. 5B and S4). For example, meiocytes expressed several histone subunits (expressed in AR cell S phase) together with microtubule-binding proteins (expressed in AR cell G2/M phase), and expressed combinations of cyclins and cyclin-dependent kinases not observed in the AR population. These results highlight many features that distinguish meiotic prophase I and any mitotic cell cycle phase in the direct predecessor cell population.

Finally, we applied our data to identify developmentally regulated genes during the AR transit amplifying stage (cells from anthers ≤1 mm in length) by calculating how strongly the expression level of each gene was correlated with (a) pseudotime and (b) the most similar cell cycle cluster. This analysis separated genes that change in expression during AR differentiation from cell cycle-regulated genes (Fig. 5D), a distinction that would have been impossible using bulk methods because of the changing proportion of cells in different phases of the cycle cell (Fig. 5C and S11). For instance, many cyclins were selectively expressed at specific cell cycle phases (Fig. 5E), as expected given their role as key cell cycle regulators. However, a subset were not strongly correlated with any cell cycle phase but rather showed stage-dependent expression regulation. Two cyclins were upregulated during the MLT, including *CycSDS-1*, one of 2 homologs of an established regulator of meiotic prophase I in *Arabidopsis*^20^. In contrast, *CycD2;1* and *CycD2;3* were constitutively expressed during the AR stage and then downregulated during the mitosis-to-meiosis transition. These genes may enable the mitotic divisions during the transit amplifying stage before the switch to meiosis.

## Discussion

Single-cell RNA-seq provides a powerful tool to elucidate cell differentiation. This is especially true in plants, because it can define cellular intermediates without requiring pre-defined marker genes, which are unknown for many important pathways. We applied scRNA-seq to pre-meiotic and early meiotic cells in maize and identified multiple gene expression stages, including a massive reorganization of the transcriptome in mid-leptotene. Intriguingly, a recent scRNA-seq survey of mouse spermatogenesis also observed a sudden jump in gene expression near the beginning of meiosis^21^; although the timing of this transition relative to meiotic prophase stages was not determined precisely, it is possible that it is analogous to the MLT we observed here.

What is the reason for such a large shift in gene expression during mid-leptotene? Our data suggest that the MLT relates not only to the nuclear events during prophase I, but also to broad changes in cellular physiology. The basic features of the MLT – a loss of ribosomal transcripts and increase in transcripts related to membrane-bound organelles and mitochondria – lead to sufficiently large changes in the meiotic cytoplasm that they were visible by electron microscopy a half century ago^17,22^. Some of these changes might have to do with the central position of meiosis to the alteration of generations, as meiosis is *the* cell division that separates the diploid sporophyte from the haploid gametophyte. Ribosome elimination during meiotic prophase could be part of a mechanism to degrade long-lived messenger RNAs and proteins prior to the gametophytic stage (as proposed previously^17^), accelerating the switch to functions encoded by the haploid genome. Future studies could address whether some of the newly synthesized mRNAs that appear during the MLT are stored for translation after meiosis. Sequestration of mRNAs is a well-established mechanism for delaying translation in animal oocytes^23^.

The MLT also has implications for the establishment of meiotic recombination hot-spots. Gene expression levels and recombination hot-spots are each shaped by similar genomic features^24^: both preferentially occur in regions of open chromatin (often in gene promoters), are associated with specific histone modifications, and are positively and negatively regulated by TF binding. Because we observed a >2 fold change in over a quarter of the transcriptome during mid-leptotene, our data imply that the genomic features modulating both gene expression and recombination can be substantially remodeled on a time-scale shorter than a substage of meiotic prophase I. This raises the possibility that every step of recombination – from double-strand break production to strand invasion to crossover interference – might occur under a different regulatory landscape, allowing for fine-tuned and independent control of each recombination step.

While cellular differentiation has often been described as the passage through a series of discrete intermediates, it is often unclear how discrete or continuous these developmental programs might be. We found both relatively continuous and discrete periods of differentiation during early germinal development in maize. So far, the single-cell field has largely focused on the important problem of cell fate decisions and bifurcations^25^, but how is the timing and rate of cellular differentiation controlled after a lineage decision has been reached? Do different types of regulatory networks and feedback loops give rise to continuous changes in gene expression vs more discrete (i.e. rapid) changes? Are discrete transitions more often under tight regulation, serving as developmental check-points? The ability to quantitatively assess the sharpness of cell state transitions using single-cell genomics provides a valuable new avenue to address these questions.

## Acknowledgements

We thank Blake Meyers and members of the Walbot lab for valuable input throughout the project. We thank the Carnegie Institution Plant Biology Imaging Facility for use of the SP8 confocal microscope and Heather Cartwright for microscope training and advice. We thank Max Staller and Blaine Marchant for critical reading of the manuscript.

## Author contributions

B.N. and V.W. conceived and designed the study. B.N. conducted experiments and analyzed the data, with input from V.W. B.N. and V.W. prepared the manuscript.

## Data availability

Raw sequencing data and processed transcript counts were deposited to NCBI GEO (accession number pending).

## MATERIALS AND METHODS

### Experimental methods

#### Plant material

Maize (*Zea mays*) inbred line W23 *bz2* was grown in a greenhouse at Stanford, CA with a 14-h day / 10-h night light cycle. For single cell RNA-seq experiments, plants were harvested between 3:30 pm – 4:30 pm (9.5 to 10.5 h into the day cycle) and processed immediately to reduce variability caused for circadian changes in gene expression.

#### Tissue dissociation

All tissue handling procedures were miniaturized so that cells could be isolated from a single floret. Anthers were dissected from the middle third of the central tassel spike and anther length was measured using a stage micrometer (Fisher Scientific). Anthers from a single upper floret were cut with a #11 scalpel every 0.5 mm and placed on a microscope slide in a 20 μL drop of anther protoplasting buffer (10 mM MES pH 5.7, 400 mM Trehalose, 2 mM CaCl_2_, 10 mM KCl, 0.1% BSA) with protoplasting enzymes: 1.25% w/v Cellulase-RS, 0.5% Macerozyme R10, 0.5% Hemicellulose, and 0.5% Pectolyase Y23 (Sigma-Aldrich catalog numbers C0615, P2401, H2125, and P5936, respectively). A second microscope slide was placed on top to minimize evaporation, using a thin piece of tape on both sides and two #1.5 coverslips to provide sufficient space between the slides to avoid crushing the anthers. Slides were incubated at 31 °C for 50 minutes, and then coverslip spacers were removed and shear force was applied by moving the top microscope slide back and forth 5-6 times to release anther protoplasts (space was still maintained between the slides by the thin pieces of tape). All chemicals were obtained from Sigma-Aldrich. The use of trehalose as an osmoprotectant instead of mannitol was important for miniaturization; this is because mannitol was prone to crystallization at the edge of small liquid droplets, while trehalose was not. Furthermore, trehalose is compatible with all downstream enzymatic steps.

#### Cell isolation

Pre-meiotic and early meiotic protoplasts were identified by their large size, picked up with a blunted 29 gauge insulin needle (BD Biosciences), and washed twice in protoplasting buffer without CaCl2. Single washed protoplasts were then picked up with a blunted 33 gauge syringe needle (Sigma-Aldrich CAD4112) and placed on the cap of a 8-tube PCR strip (Axygen Low Profile 8-Strip PCR Tubes; Fisher Scientific 14-223-505). The presence of an isolated single cell without attached debris was confirmed microscopically (10X magnification, Nikon Diaphot) during both cell pick-up and release. Each isolated cell was photographed, providing both an image with which to measure cell size and a time stamp for when each cell was isolated. Based on the diameter of the droplets visible in these images, we estimate that cells were isolated in a typical buffer volume of 10-50 pL. After isolating each set of 8 cells, the PCR caps were attached to PCR tubes and flash frozen in liquid nitrogen. Frozen cells were stored at −80 °C.

#### Illumina library preparation

Sequencing libraries were prepared using a protocol based on CEL-seq2^26^, with three modifications: (i) the lysis step was modified to enable split-cell technical replicates, (ii) comparable but different enzymes (e.g. different companies) were used, either for convenience or because the enzymes were found to be more efficient with maize RNA, and (iii) the unique molecular identifying (UMI) length was increased from 6 to 10 nucleotides. To avoid correlations between batch effects and biological information, sample locations and pooling schemes were randomized at several points during library synthesis (Table S1).

Briefly, frozen cells were thawed and immediately lysed by adding 2.1 μL lysis buffer (0.1% Triton X-100, 1.5 mM dNTP mix, 1:315,000 Ambion ERCC spike in controls) and incubating at 65 °C for 2 minutes. Then 1 μL (half) of the cell lysate was transferred to a fresh tube and 1 μL 1.5 μM barcoded oligo(dT) primer was added to each split-cell sample. Samples were incubated at 65 °C for 3 minutes then placed on ice. Reverse transcription was initiated by adding 1 μL of Life Technologies SuperScript IV reverse transcription mix (4:1:1:1 ratio of 5x Superscript IV buffer, RnaseOUT, Superscript IV, 100 mM DTT) and incubated at 42 °C for 2 minutes, 50 °C for 10 minutes, 55 °C for 5 minutes, and 70 °C for 10 minutes. Excess primers were digested with exonuclease 1 (New England Biolabs).

Next, second strand synthesis was performed using the NEBNext mRNA Second Strand Synthesis Module (New England Biolabs). Sets of 16 individually barcoded split-cell samples were pooled and DNA was purified using a 1.2 ratio of AMPure XP beads (Beckman Coulter). Samples were then amplified with the MEGAscript T7 *in vitro* transcription kit (Thermo Fisher Scientific) at 16 °C overnight. Amplified RNA was purified with RNAClean XP beads (Beckman Coulter) and a second round of reverse transcription was performed with SuperScript II reverse transcriptase (Life Technologies) using a tagged random hexamer primer (5’-GCCTTGGCACCCGAGAATTCCANNNNNN). Libraries were then amplified with 10-15 rounds of PCR using the NEBNext Ultra II Q5 Master Mix (New England Biolabs) and Illumina primers, and libraries were purified and size selected with AMPure XP beads. Library quantity and size distribution was assessed with an Agilent Biolanalyzer High Sensitivity DNA kit.

Libraries were sequenced on an Illumina HiSeq 2500 instrument at the University of Delaware Sequencing and Genotyping Center with paired-end 76 bp reads. Primer sequences can be found in Table S5. All primers were synthesized by phosphoramidite chemistry at the Stanford Protein and Nucleic Acid Facility (Stanford, CA).

#### Quantitative PCR (qPCR)

Two of 3 anthers from a floret were placed on a cleaned microscope slide in 10 μL water. Anthers were then cut in half with a #11 scalpel and meiotic cells were gently squeezed out. Remaining anther material was removed, and meiotic cells with excess water were placed in a PCR tube (Axygen) and flash frozen in liquid nitrogen. Extruded meiocytes were lysed with 0.1% Triton X-100 (Thermo Fisher Scientific) and reverse transcription was performed without further purification using SuperScript IV reverse transcriptase (Life Technologies) and oligo(dT)12-18 primer (Thermo Fisher). Quantitative PCR was performed using TaqMan primers synthesized by Integrated DNA Technologies (Table S6) on a BioRad CFX instrument. Three to 4 replicates were performed per condition, then cycle threshold (Ct) values for replicate samples were averaged and the mean Ct of the reference gene (*Acp2*) was subtracted.

#### Cytological staging and image acquisition

One of 3 anthers from a floret was fixed in 4% paraformaldehyde at room temperature for 2 h. Anthers were washed and then fixed meiotic cells were extruded in Prolong Diamond antifade media (Life Technologies) with 10 μg/mL Hoechst 33342 (Sigma-Aldrich). Finally, a #1.5 coverslip (Zeiss) was placed on top. Confocal image stacks were taken with a Leica SP8 microscope using a 63X 1.4 n.a. oil immersion objective, a 405 nm excitation laser, an AOTF emmission filter set to capture light between 495 nm and 562 nm, and a Nyquist resolution voxel size of 58.2 nm by 58.2 nm by 299 nm. All images were processed uniformly in ImageJ^27^ Fiji^28^ by applying rolling ball background subtraction with a 50 pixel radius and then linearly scaling the maximum and minimum intensity values.

Cytological stage was scored using established criteria^11^: (i) premeiotic interphase / early leptotene – centrally located nucleolus, round knobs, chromatin threads not visible or visible as localized patches; (ii) leptotene stage 1 – centrally located nucleolus, round knobs, chromatin threads condensed throughout the nuclear volume; (iii) leptotene stage 2 – same as leptotene stage 1 except the nucleolus was located at the nuclear periphery; (iv) prezygotene – same as leptotene stage 2 except the knobs were clearly elongated. Cytological scoring was performed blinded to sample identity (i.e. without any knowledge of the anther length or qPCR staging).

### Computational methods

#### Library mapping

One of each paired-end sequencing read contained transcript sequence; these reads were mapped to the B73 reference genome (AGPv4) using Hisat2^29^. The second read pair contained cell barcodes and a 10 bp unique molecular identifier^30^ (UMI). For transcript counting, read-pairs that mapped to the same gene with one or fewer bp differences in their UMIs were counted as originating from a single first-strand cDNA molecule. Allowing for a 1 base error in the UMIs reduced overcounting artifacts resulting from sequence read errors.

During library mapping, two errors were detected in the cell barcode reverse transcription primers. First, primer ‘CelSeq_dT-1s’ had an extra nucleotide added to the cell barcode (Table S5), which was apparently the result of a systematic primer synthesis error; the cell barcode was updated to account for this synthesis error and no data were lost. Second, very few reads mapped to the primer ‘CelSeq_dT-4s’, which could be attributed to a second primer synthesis error or a mistake during primer dilution; samples with primer dT-4s were lost from the dataset, and so a subset of cells do not have technical replicates (16 cells out of the 144 that passed QC criteria).

#### Quality control

Before applying quality control (QC) criteria, data from split-cell technical replicates were pooled together. Pooling split-cell replicates for QC was important because technical replicates were later used to assess analysis quality, and so blinding QC to replicate information avoided selecting for cells where the technical replicates were in agreement. For QC, cells were excluded if under 71.8% of mapped reads mapped to the 10 maize chromosomes; the 71.8% cutoff was chosen because it is 1.5 median absolute deviations from the population median (Fig. S1A). This QC criterion excluded cells with a high proportion of reads mapping to the ERCC spike-in controls and/or the organellar (mitochondria/chloroplast) genomes. Four additional cells were excluded because their transcript abundance was poorly correlated with every other cell in the dataset (Pearson’s correlation < 0.6; Fig. S1B). These quality control criteria did not substantially enrich for cells at any given anther length.

#### Calculation of pseudotime

Genes with a low transcript abundance (<100 UMIs across all samples) were excluded from analysis; 12,902 genes passed this abundance criteria. In addition, 375 genes that appeared to be expressed at specific cell cycle phases (see below) were excluded. The resulting data (12,527 genes by 272 split-cell samples) were normalized by dividing by the total number of transcripts in each cell and multiplying by one million (transcripts per million normalization). Then the dataset was log transformed after adding a pseudocount of 11, corresponding to 0.5 UMIs per cell on average. To calculate pseudotime, the top 2000 most variable genes were selected, the data were transformed by principal component analysis (PCA), and a principal curve was fit to the top 10 PCs using the R package princurve^9^.

#### Calculation of pseudotime velocity

Pseudotime velocity was calculated repeatedly after drawing a subset of samples from the complete dataset (bootstrapping), which served to both smooth out noise in the calculation and provided a way to assess uncertainty in the velocity estimate. For each of 1000 bootstrap rounds, 24 samples were randomly drawn from each of 7 anther size bins (Fig. S5A). Binning the cells by anther size enforced even sampling across developmental time. Pseudotime was then calculated using the principal curves method. Each step of the pseudotime calculation was repeated for every bootstrap sample, including selecting the top 2000 most variable genes, PCA, and the principal curve fit. Then samples were ordered by increasing pseudotime, and pseudotime velocity was calculated as the slope of a linear least squares fit to pseudotime rank vs pseudotime with a rolling window size of 10 samples (see Fig. 2B). Ninety-five percent confidence intervals were calculated as the 0.025 and 0.975 quantiles of the bootstrap values. Estimates of pseudotime velocity were robust to the binning scheme for bootstrapping (Fig. S5B) and the window size selected for the linear slope calculations (Fig. S5C).

To link pseudotime velocity to estimated developmental time (e.g. bottom legend in Fig. 2C), the mean pseudotime rank of each sample was first calculated from the bootstrap estimates. Each sample was then attributed a developmental time so as to preserve (i) the rank order estimated from pseudotime and (ii) the total actual time (168 hours) represented by the single-cell time-course. Estimates of developmental time recovered the expected relationship with the age of each anther, as a linear fit of estimated developmental time (in hours) vs estimated anther age (in days, based on anther length) had a slope of 24.1 ± 0.6 (mean ± standard error; R ‘lm’ function).

#### Comparison between pseudotime velocity and clustering

Eight single-cell RNA-seq clustering methods were applied to identify cell stages using the log-transformed dataset after selecting the 2000 most-variable genes. SC3^31^, RaceID3^32^, SNN-cliq^33^, and SEURAT^34^ were applied using the published R or python packages with standard parameters. SINCERA^35^ was applied using Z-transformed data and the ‘graph’ clustering method. PCAreduce^33^ was applied using the largest probability method (method = ‘M’). Hierarchical clustering was performed using base R functions and the ‘ward.D’ clustering method. PCA + k means was performed using the R function ‘kmeans’ applied to the first two principal components of the PCA-transformed data.

SEURAT, SINCERA, and SNN-cliq provided a single clustering solution. The other 5 methods required the selection of a predefined number of clusters. These methods were applied with a range of cluster numbers from 2 to 5, and the largest number of clusters that correctly grouped at least 90% of the split-cell technical replicates was selected. The performance of all methods as a function of cluster number (k) can be seen in Fig. S6B (assessed based on the percentage of split-cell replicates grouped together). A consensus clustering solution was also considered by plotting a heatmap of how often every method grouped the same cells together (Fig. S6A), similar to the approach taken by SC3^31^.

Pseudotime velocity was used to identify cell stages by selecting each peak in pseudotime velocity as a potential stage boundary. A range of stage numbers was considered (k = 2 to 5) by ranking each stage boundary based on the height of the corresponding pseudotime velocity peak; for example, the 2 stage solution (k = 2) for pseudotime velocity was provided by splitting the samples at the position of the largest velocity peak, the 3 cluster solution (k = 3) was selected by splitting the samples at the largest and second largest velocity peaks, etc. Pseudotime velocity performed better than all clustering methods at grouping split-cell technical replicates for every value of k (Fig. S6B).

#### Estimating the fraction of genes that change in expression during the MLT

Transcript counts for split-cell technical replicates were pooled (to provide a single value for each gene and each cell) and then the data were TPM-normalized and log-transformed as described in the ‘calculation of pseudotime’ section (above). Next, the mean log_2_ expression level of each gene was calculated for ‘early AR cells’ (cells with pseudotime ≤ 10, a total of the 20 cells), M1 stage meiotic cells (orange in Fig. 2C), and M3 stage meiotic cells (dark red in Fig. 2C). The log_2_ fold change in expression during the MLT (FC_MLT_) was calculated as the difference between the mean log_2_ expression in M3 stage cells and M1 stage cells (the stages just before and just after the MLT), and the log_2_ fold change in expression from early AR to M1 (FC_AR>M1_) was calculated similarly.

To quantify the fraction of genes that change >2 fold during the MLT, the log_2_ fold-change values for each gene, 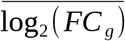, were modeled as a response variable with homoscedastic, normally distributed measurement errors:

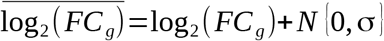

Under this noise model, the population estimate for the fraction of genes with a log_2_(FC) exceeding any given threshold becomes a deconvolution problem. Deconvolved estimates of the fraction of genes with a log_2_(FC) greater than 1 or less than −1 were calculated using the ‘cdf.decon’ function of the R package ‘spsurvey’^36^. This calculation was repeated for 1000 bootstrap samples using the R package ‘boot’^37^, and bias-adjusted estimates ±95% confidence intervals were reported.

#### Identifying differentially expressed genes

For each gene, a cubic smoothing spline with 5 degrees of freedom was fit to the curve of pseudotime vs expression level. An example of these smooth splines can be seen in Fig. 2D (black lines on the scatter plots to the right). Then the proportion of variance explained by the spline was calculated as:

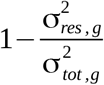

where 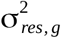 is the variance of the residual expression levels for gene *g* after subtracting the smooth spline, and 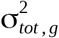 is the variance in the original data for gene *g*. The variance explained by the spline was then used as a test statistic for each gene, and a *p*-value was calculated from this statistic using a permutation test. *p*-values were adjusted for multiple hypothesis testing with Holm’s method.

#### Gene clustering, pathway enrichment analysis, and gene family identification

All differentially expressed genes were grouped by kmeans clustering. A range of cluster numbers (from 2 to 15) was considered and k = 6 was selected because it was the largest k that returned clearly distinguishable clusters. GO term enrichment was calculated for each cluster with the AgriGO^15^ web tool, using the *Zea mays* landing page and default parameters. PlantTFDB^38^ was used to define which differentially expressed genes were transcription factors.

#### Identifying cell cycle-regulated genes and cell cycle phases

To find cell cycle regulated genes, we first took advantage of our split-cell technical replicates and identified transcripts with significant cell-to-cell variability after controlling for developmental progress. A permutation-test ANOVA was performed to determine which transcripts had significantly greater variation between cells isolated from the same floret than can be explained by technical variability. Of the 182 genes with significant cell-to-cell variability, the majority (74%) formed clusters readily attributable to phase in the cell cycle. These initial cell cycle clusters were expanded to include other genes with a correlated expression pattern (Pearson’s correlation ≥ 0.4) in cells isolated from ≤ 1 mm anthers (the transit amplifying stage). A total of 375 cell cycle-regulated genes was identified (Table S2).

By grouping the cell cycle-regulated genes using hierarchical clustering (Fig. S4), each gene was assigned to a specific cell cycle phase. For instance, one cluster of genes contained 35 histone subunits (43% of the 81 genes in the cluster). Histone transcripts are upregulated during S-phase in most eukaryotes, and so this cluster was assigned to S-phase. The inferred order of expression of each cell cycle gene cluster based on principal component analysis (Fig. 5A) or hierarchical clustering (Fig. S4) reproduced the known order of the associated cell cycle phases during mitosis. This provides the first genome-wide, experimentally determined list of cell cycle-regulated genes in plants without the use of chemical inhibitors for synchronization. Each cell was assigned to a cell cycle phase by hierarchical clustering – i.e. clustering all cells based on the expression of the 375 cell cycle-regulated genes (Fig. S4B).

It should be pointed out that a gene cluster could be associated with every phase of the mitotic cell cycle except for G1. Putative G1 genes were defined as those genes that were both (i) negatively correlated with the cumulative expression of all 375 cell cycle genes (Pearson’s correlation ≤ −0.3) in cells isolated from ≤ 1 mm anthers and (ii) not strongly correlated with genes from any individual cell cycle phase (Pearson’s correlation ≤ 0.1 for the most similar cell cycle cluster). There was far less confidence in the G1 gene assignments compared to the 375 genes associated with every other cell cycle phase. These ‘G1 genes’ were included in Fig. 5A and 5B for visualization purposes and were reported in Table S2, but were not considered as ‘cell cycle-regulated’ genes for any other analyses.

For Fig. 5D and Fig. 5E, cyclins were defined as genes that encode proteins with a cyclin N-terminal domain (Interpro ID IPR006671).

## List of supplementary materials (available online)

**Fig. S1.** Histogram of the number of cells collected at each anther size.

**Fig. S2.** Quality control criteria.

**Fig. S3.** Reproducibility of technical replicates.

**Fig. S4.** Cell-cycle regulated gene clusters.

**Fig. S5.** Evaluation of bootstrap sampling and parameter choice for pseudotime velocity calculations.

**Fig. S6.** Comparison of stages determined by pseudotime velocity with single-cell clustering methods.

**Fig. S7.** Expression of marker genes for the mid-leptotene transition (MLT).

**Fig. S8.** Effect of normalization on the estimated percentage of genes that change ≥2 fold during the MLT.

**Fig. S9.** Assessment of transcript diversity at each cell stage

**Fig. S10.** Dynamic regulation of genes linked to sub-cellular organelles during early germinal development.

**Fig. S11.** MA plot of simulated bulk data depicting gene expression changes between cells from 0.5 mm and 1.0 mm anthers.

**Table S1.** Sample information for single isolated AR and PMCs.

**Table S2.** Cell cycle-regulated gene clusters.

**Table S3.** Differentially expressed genes during early germinal development in maize anthers.

**Table S4.** Enriched GO terms.

**Table S5.** Primer sequences for scRNA-seq library construction.

**Table S6.** Primer sequences for quantitative PCR.

